# Digestive parameters during gestation of Holstein heifers

**DOI:** 10.1101/2020.04.20.050773

**Authors:** Jéssica Marcela Vieira Pereira, Marcos Inácio Marcondes, Sebastião de Campos Valadares Filho, Joel Caton, Anna Luiza Lacerda Sguizzato, Alex Lopes Silva, Julia Travassos da Silva, Valber Carlos Lima Moraes, Lucas Faria Gomes, Polyana Pizzi Rotta

**Author notes:** Corresponding author (PPR). These authors contributed equally to this work (Jéssica Marcela Vieira Pereira, Marcos Inácio Marcondes, Joel Caton, Polyana Pizzi Rotta). These authors also contributed equally to this work (Sebastião de Campos Valadares Filho, Anna Luiza Lacerda Sguizzato, Alex Lopes Silva, Julia Travassos da Silva, Valber Carlos Lima Moraes, Lucas Faria Gomes).

## Abstract

Our objective was to estimate nutrient intake, ruminal flow, total apparent and ruminal digestibility, rates of passage and digestion, ruminal and omasal pH, blood metabolite concentrations, and body measurements during gestation of Holstein heifers. Eleven pregnant Holstein heifers, 8 of which fitted with a rumen cannula (450 ± 27.6 kg of body weight and 20 ± 3.5 months of initial age) were used. All heifers received the same diet composed of corn silage, soybean meal, corn meal, minerals and vitamins, with a corn silage:concentrate ratio of 50:50 (on a dry matter basis), aiming an average daily gain of approximately 1.0 kg. The sampling periods were established according to the days of gestation: 145, 200, and 255 with a duration of 10 days per period. Total fecal samples were collected to estimate dry matter, organic matter, crude protein, and neutral detergent fiber digestibility. Blood samples were collected to analyze metabolites (non-esterified fatty acids, β-hydroxybutyrate, urea, and glucose). Data were analyzed as a repeated measurements scheme, using MIXED procedure, with differences declared when *P* < 0.05. Dry matter intake expressed in kg/day increased from d-145 to d-200, and remaining stable until d-255 of gestation. The same results were observed for organic matter and crude protein intake, increasing 15.0 and 35.8% respectively. In contrast, when dry matter intake was evaluated as % body weight, we observed a decrease of 16.7% from d-200 to d-255. Days of gestation did not influence ruminal flow of dry matter, organic matter, crude protein, and neutral detergent fiber. We observed an increase in the ruminal digestibility of neutral detergent fiber by 20.5%. The apparent total-tract digestibility of dry matter, organic matter, and crude protein changed over days of gestation, with an increase of 11.9, 8.5, and 9.8%, respectively, when comparing d-145 with d-200. The rate of digestion of neutral detergent fiber increased from 2.0 to 3.5% h^-1^. Glucose levels decreased, while β-hydroxybutirate and non-esterified fatty acids increased from d-145 to d-255. In conclusion, our results demonstrate a reduction in dry matter intake in % body weight due to pregnancy. It also shows an increase in total apparent digestibility through gestation, which imply a greater efficiency of use of nutrients by pregnant animals. Thus, further research is still needed to consolidate such results and to elucidate the mechanism about nutrient usage during the final third of gestation in heifers.

## Introduction

According to the Hoffman [1], heifers should have the first parturition with at least 94% of their mature body weight, considering the fetal weight. Thus, their nutritional plan requires strategies to maintain constant and satisfactory gains until the end of gestation [2]. According to Mohd [3] the advances in reproduction were superior to the nutritional advances of heifers, producing precocious animals that may present low milk yield in the first lactation.

Literature lacks information on digestive aspects of ruminal flow, intake, passage and digestion rates of pregnant replacement heifers. However, several studies addressing nutrient fluxes of non-pregnant lactating animals are found in the literature [4, 5]. Eventually, mathematical models were developed to increase feed efficiency and to estimate rates of passage, digestion, and to estimate outflow of rumen digesta [6, 7, 8], however even these models were not developed or evaluated with pregnant heifers.

Data regarding nutrition of dairy cattle during gestation are focused on lactating dairy cows and are scarce for heifers [2, 9, 10]. It is well known that dry matter intake decreases during the final third of gestation due to the greater size of the gravid uterus and the increased estrogen concentration [11, 12]. This decrease in dry matter intake may reach up to 50% in the last 2 months of gestation in dairy cows [12]. However, it is not known whether dry matter intake in heifers present a similar decrease since they are still growing; such information would be of great use to improve management practices in pregnant heifers.

The hypothesis of this study was that the digestive parameters of pregnant heifers would be different from data in the literature for pregnant cows and would also be not altered during pregnancy. Thus, the objectives of this study were to estimate ruminal flow, total and partial digestibility of dry matter and diet constituents, intake, passage and digestion rates, ruminal and omasal pH, blood metabolite concentrations, and body measurements during gestation in Holstein heifers.

## Materials and methods

### Animals and management

All procedures were previously approved by the Animal Ethics and Welfare Committee of the Federal University of Viçosa according to protocol number 020-2016. Eleven pregnant Holstein heifers, 8 of them fitted with a rumen cannula, with a mean body weight of 450 ± 27.6 kg and mean initial age of 20 ± 3.5 months were used in this experiment. The sampling periods were established according to the days of gestation: 145, 200, and 255 ± 4.45, with duration of 10 days for each period. The animals were allocated to individual stalls of 10 m^2^. Prior to the start of the experiment, heifers underwent a 30-day adaptation period to experimental facilities and conditions. Heifers received the same diet (Table 1) for an average daily gain of 0.98 ± 0.30 kg/day, according to NRC (2001).

**Table 1.** Ingredients and chemical composition of feed used in the experimental diet.

| Item | % of DM |
| --- | --- |
| Ingredients |  |
| Corn silage | 50.0 |
| Soybean meal | 25.0 |
| Ground corn | 21.6 |
| Limestone | 1.07 |
| Sodium bicarbonate | 0.75 |
| Magnesium oxide | 0.25 |
| Salt | 0.50 |
| Vitamins | 0.76 |
| Mineral mix<sup>1</sup> | 0.02 |
| Chemical composition |  |
| Dry matter | 56.3 |
| Organic matter | 95.2 |
| Crude protein | 17.7 |
| Neutral detergent fiber | 32.2 |
| Ether extract | 2.7 |
| Non fiber carbohydrates | 42.6 |
| Indigestible neutral detergent fiber | 10.3 |

<sup>1</sup>Mineral mix composition = 0.83 mg/kg of cobalt sulfate, 38.6 mg/kg of copper sulphate, 0.215 g/kg iron sulphate, 2.39 mg/kg of sodium selenite, 0.167 g/kg of zinc sulfate, and 1.71 mg/kg potassium iodate.

The roughage:concentrate ratio was 50:50 (dry matter basis) and was delivered daily at 0700 and 1500 h. Animals were allowed 5% of orts on fresh basis. Once a week, corn silage was sampled and dried in a ventilated oven at 55°C for 72 h to adjust the diet dry matter content.

### Intake and digestibility trial

Dry matter intake was daily measured by weighing offered and refused feedstuffs. To estimate nutrients digestibility, total feces were collected from all heifers for 3 consecutive days in first period: 145-148 days of gestation, second period: 200-203 days of gestation, and third period 255-258 days of gestation. After 24 h of collection, the total feces were weighed and approximately 250 g was oven-dried (55°C for 72 h). After each collection period, a proportional sample was composed using samples and data from the 3 days of fecal collection, based on fecal dry matter production of each day. Then, feces and feed samples were ground using a mill (Willye, model TE-680, TECNAL, Brazil) with a 1- and 2-mm sieves and stored for further chemical analyses.

### Omasal and ruminal sampling

A continuous infusion of 5 g/day of Co-EDTA (0.7 g cobalt/day) from the 1^st^ to the 6^th^ day of collection was performed during each period (first period: 145-150 days of gestation, second period: 200-205 days of gestation, and third period 255-260 days of gestation) using 2 peristaltic pumps (model BP-600.4, Colombo, Paraná, Brazil). A total of 8 samples per animal were collected from the omasum at a 9-h interval for 3 days; sampling times were 0200, 0500, 0800, 1100, 1400, 1700, 2000, and 2300 h. The technique, that was developed by Huhtanen [13] and adapted by Leão [14], was used for sampling of the omasal digesta, where approximately 250 mL of digesta was used for a bacterial isolate and 500 mL was used for the ruminal flow estimation [15]. The samples were stored in plastic containers at -20 °C for further analysis.

At the end of each experimental period, samples of omasal digesta were thawed at room temperature and one composite was created per animal, following the recommendations of Rotta [15]. Thus, 4 L were obtained to estimate the omasal flow, which was filtered using a 100 μm nylon filter with an open area of 44% (Sefar Nytex 100/44; Sefar, Thal, Switzerland), thereby obtaining two phases: liquid particles and solid particles. Thus, the double marker system technique was used to estimate the digesta omasal flow, using cobalt and indigestible neutral detergent fiber as markers. All samples were stored in plastic containers and kept at -80° C for further lyophilization (LP510; Liobras, São Paulo, Brazil). After lyophilization, samples were milled at 2- and 1-mm sieves mills (Willye, model TE-048, TECNAL, Brazil).

Ruminal contents were sampled during the same times as omasal sampling, resulting in nine samples. Samples were manually collected at the liquid-solid interface of the rumen, filtered through a 100 μm nylon filter (Sefar Nitex; Sefar, Thal, Switzerland; porosity of 50 μm), and were subjected to pH measurement by using a digital potentiometer (pH-221, Lutron Electronics, Taiwan).

On the 7 th day of collection of each experimental period, total rumen emptying was carried out 4 h after feeding to estimate the rate of passage and digestion [16]. After removal of all ruminal contents, the digesta was weighed and filtered in a double layer of cheesecloth for separation of solid and liquid fractions, which were then weighed and sampled. After sampling, the digesta was reconstituted and returned to the rumen of the respective animals. On the 9 th day, the entire rumen emptying procedure was repeated before morning feeding. The collected samples were lyophilized and ground in a knife mill with 1 mm sieve (Willye, model TE-680, TECNAL, Brazil). Thus, composite samples of the dried solid and liquid fractions after emptying the rumen were obtained and were based on the dry weight of each sample.

### Biometric Measurements

At the end of each experimental period (first period: 154 days of gestation, second period: 209 days of gestation, and third period 264 days of gestation), body weight, withers height, croup height, body length, and thoracic perimeter were measured while the animals were kept in a standing position within the chute.

### Blood measurements

Blood samples were collected on the 10 th day of each period. The samples were collected via jugular venipuncture by using vacuum tubes with clot activator and gel for serum separation (BD Vacutainer® SST® II Advance®, São Paulo, Brazil) in order to quantify urea, nonesterified fatty acids, and β-hidroxibutirate; an extra tube with EDTA and sodium fluoride (BD Vacutainer® Fluorinated/EDTA, São Paulo, Brazil) was used to quantity plasma concentration of glucose. After the collection, samples were centrifuged at 3600 × g for 20 min, and serum and plasma were immediately frozen at - 20°C until further analysis.

Bioclin® kits were used to quantity urea (K056) and glucose (K082), nonesterified fatty acids, and β-hidroxibutirate were analyzed by using Randox® kits (FA115 and RB1007, Antrim, United Kingdom). All mentioned analyses were conducted using an automated biochemical analyzer (Mindray, BS200E, Shenzhen, China).

### Laboratory analysis and calculations

Samples of omasal digesta, feces, feed, and ruminal emptying were analyzed for dry matter, organic matter, and nitrogen ([17]; method 934.01 for dry matter, 930.05 for organic matter, and 981.10 for nitrogen, respectively). The ether extract analysis was performed according to AOAC recommendations ([18]; method 945.16). The neutral detergent fiber was analyzed, without the addition of sodium sulphite, but with the addition of detergent thermostable alpha amylase ([19]; INCT-CA method F-001/1). Ash correction was performed in the neutral detergent fiber residues ([19]; method 942.05). Indigestible neutral detergent fiber was estimated according with Valente [20], in triplicate for omasal digesta (particle phase), fecal samples, and rumen emptying samples. Non-fibrous carbohydrates were calculated according to Van Soest [21], with no correction for urea.

The total-tract apparent digestibility was calculated as:

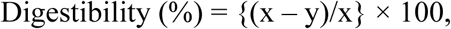

where x (kg/day) and y (kg/day) are the intake and the output in feces of each component, respectively.

Ruminal digestibility coefficients were estimated by measuring the difference between the rate of nutrient intake and the flow of the nutrients through the rumen. The calculation of intestinal digestibility was estimated by the concentration of nutrients in omasal digesta and feces. The flow of the omasal digesta was estimated through the reconstitution of digesta technique [22] using the double marker system. Cobalt was used as the liquid phase marker and the indigestible neutral detergent fiber as the particle phase marker. The reconstitution factor was calculated based on the concentrations of the markers during the different phases of the digesta [23].

The rates of ingestion (k_i_), passage (k_p_), and digestion (k_d_) were calculated using the pool-and-flux method [16], according to the following models:

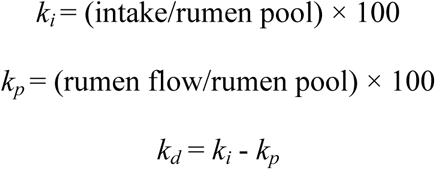

Where, *k*_*i*_ = ingestion rate of feed fractions (%/h); intake = feed intake (kg/h); rumen pool = amount of total rumen dry matter (kg); *k*_*p*_ = passage rate of feed fractions (%/h); rumen flow = amount of dry matter or nutrients in the omasum (kg/h); *k*_*d*_ = digestion rate of diet fractions (%/h).

### Statistical analysis

Data were analyzed as a repeated measurements scheme using the SAS MIXED procedure (SAS Institute Inc., version 9.4) according to the statistical model:

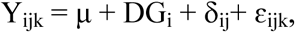

where Y_ijk_ = dependent variable, μ is the overall mean, DG_i_ = fixed effect of days of gestation, δ _ij_ the random error with mean of 0 and a variance σ ^2^ δ, the variance between the days of gestation is equal covariance between repeated measures within animals and ε_ijk_ is the model error with a mean of 0 and a variance of τ^2^. Days of gestation was considered as repeated measures, and data from ruminal and omasal pH measurements were also included in the statistical model as time-repeated measures (split-plot of DG). The fixed time effects and the interaction between time and days of gestation were tested by using the appropriate covariance structure. The following covariance matrices were tested: components of variance, composite symmetry, heterogeneous composite symmetry, and first order heterogeneous auto-regression. The selection of the covariance matrix was based on the Akaike-corrected criterion, and the composite symmetry matrix was chosen. Results are presented as least squared means and their standard error of the mean. Differences were declared when *P* < 0.05.

## Results

### Intake, ruminal flow and digestibility

The dry matter intake (kg/day) increased (*P* = 0.03) at d-200 and d-255 compared to d-145 (Table 2). As for nutrient intake, the same pattern was observed, with an increase of 14.7% (*P* < 0.01) for organic matter, 41.7% (*P* < 0.01) for crude protein, at 145 and 200 days of gestation, and for neutral detergent fiber there was no difference. Regarding to dry matter intake in % body weight (Fig 1), there was no difference (*P* > 0.05) between 145 to 200 days of gestation, but there was a 16.7% (*P* < 0.01) reduction from 200 to 255 days of gestation.

**Table 2.** Intake (n = 11), ruminal outflow (n = 8), ruminal (n = 8) and apparent total-tract digestibility (n = 11) of Holstein heifers in different days of gestation.

| Item | Day of gestation |  |  | *P*-value |
| --- | --- | --- | --- | --- |
|  | 145 | 200 | 255 |  |
| Intake, kg/day |  |  |  |  |
| Dry matter | 8.4 ± 0.48<sup>b</sup> | 9.7 ± 0.48<sup>a</sup> | 9.0 ± 0.51<sup>ab</sup> | 0.03 |
| Organic matter | 7.5 ± 0.36<sup>b</sup> | 8.6 ± 0.36<sup>a</sup> | 8.4 ± 0.38<sup>a</sup> | < 0.01 |
| Crude protein | 1.2 ± 0.76<sup>b</sup> | 1.7 ± 0.76<sup>a</sup> | 1.6 ± 0.08<sup>a</sup> | < 0.01 |
| Neutral detergent fiber | 2.8 ± 0.12 | 2.7 ± 0.12 | 2.7 ± 0.13 | 0.64 |
| Ruminal outflow, kg/day |  |  |  |  |
| Dry matter | 5.2 ± 0.28 | 5.5 ± 0.28 | 5.2 ± 0.31 | 0.54 |
| Organic matter | 4.3 ± 0.24 | 4.5 ± 0.24 | 4.3 ± 0.26 | 0.57 |
| Crude protein | 1.0 ± 0.07 | 1.1 ± 0.07 | 1.1 ± 0.08 | 0.57 |
| Neutral detergent fiber | 1.2 ± 0.14 | 1.4 ± 0.14 | 1.7 ± 0.16 | 0.08 |
| Ruminal digestibility, % |  |  |  |  |
| Dry matter | 54.3 ± 2.64 | 55.7 ± 2.64 | 55.7 ± 3.15 | 0.75 |
| Organic matter | 58.6 ± 2.37 | 58.8 ± 2.37 | 59.8 ± 2.63 | 0.89 |
| Neutral detergent fiber | 42.4 ± 4.22<sup>b</sup> | 51.1 ± 4.22<sup>ab</sup> | 62.1 ± 4.88<sup>a</sup> | 0.03 |
| Apparent total-tract digestibility, % |  |  |  |  |
| Dry matter | 72.1 ± 1.35<sup>a</sup> | 79.6 ± 1.35<sup>b</sup> | 79.9 ± 1.49<sup>b</sup> | < 0.01 |
| Organic matter | 75.0 ± 1.29<sup>a</sup> | 81.4 ± 1.29<sup>b</sup> | 82.5 ± 1.43<sup>b</sup> | < 0.01 |
| Crude protein | 75.2 ± 1.21<sup>a</sup> | 82.7 ± 1.21<sup>b</sup> | 82.7 ± 1.33<sup>b</sup> | < 0.01 |
| Neutral detergent fiber | 67.0 ± 2.35 | 70.4 ± 2.35 | 73.9 ± 2.59 | 0.15 |

<sup>a-b</sup>Means within a row with different superscripts differ ( $P < 0.05$ ).

**Fig 1.**
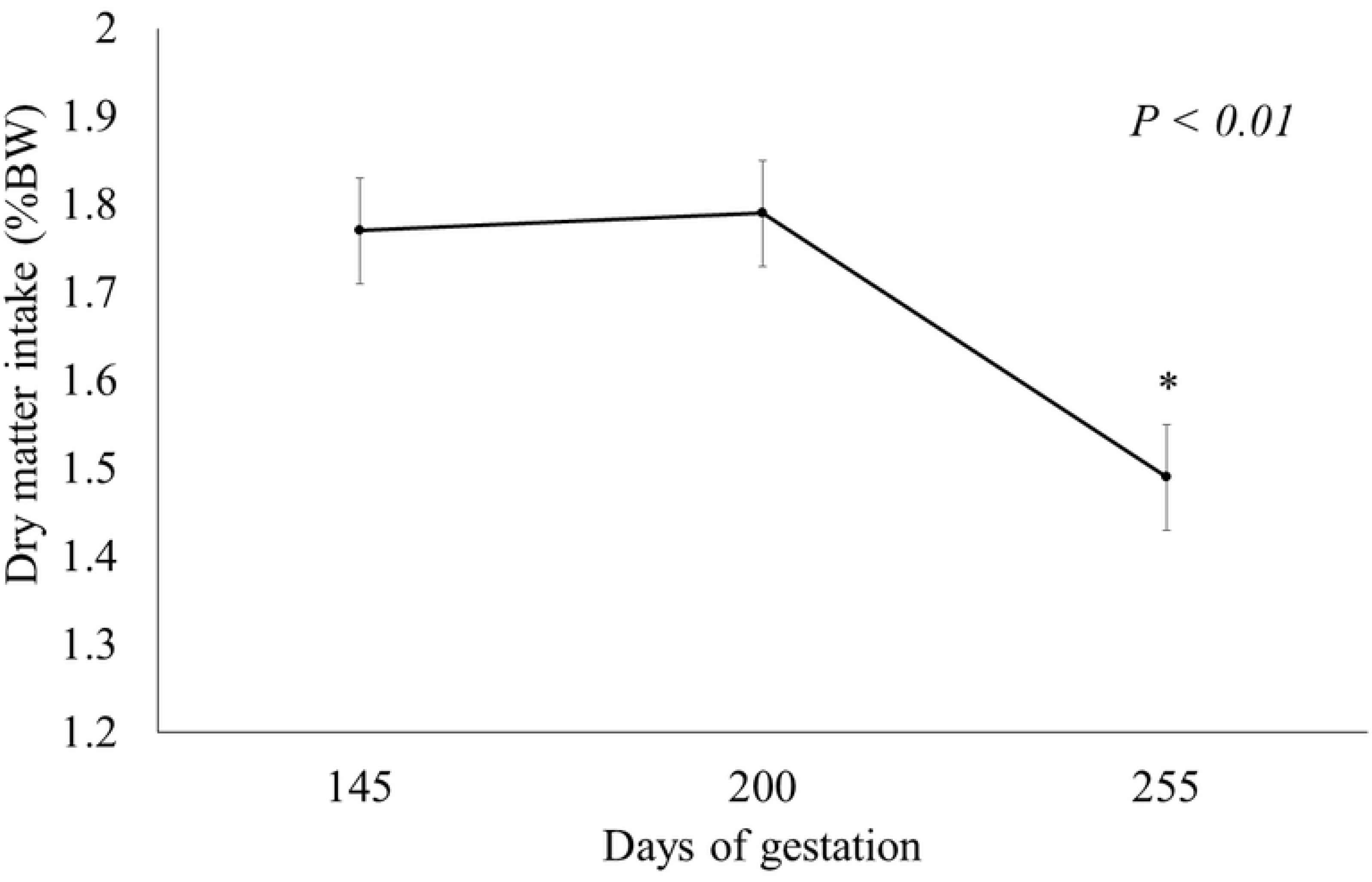
Dry matter intake (% BW) in different days of gestation of Holstein heifers (number of animals = 11). * Indicative of significance in the days of gestation (*P* < 0.05)

Ruminal flows of dry matter, organic matter, crude protein, and neutral detergent fiber were not affected by days of gestation (Table 2). The same pattern was observed for ruminal digestibility of dry matter and organic matter, but neutral detergent fiber increased by 46.6% from 145 to 255 days of gestation (*P* = 0.03). The apparent total-tract digestibility of dry matter, organic matter, and crude protein was altered (*P* < 0.01) by days of gestation, with an increase of 11.9, 8.5, and 9.8%, respectively, when comparing the 145 and 200 days of gestation. There was no difference (Table 2) for neutral detergent fiber digestibility among days of gestation.

### Digestion kinetics

Rumen pools and kinetics data are shown in Table 4. The values for dry matter *k*_*i*_, *k*_*p*_, and *k*_*d*_ did not differ among days of gestation. The means of dry matter *k*_*i*_, *k*_*p*_, and *k*_*d*_ were 8.0, 4.7, and 3.3 % h^-1^, respectively. The values for neutral detergent fiber *k*_*i*_ and *k*_*p*_ did not differ among days of gestation, and the means were 4.3 and 2.0 % h^-1^, respectively. The *k*_*d*_ of neutral detergent fiber increased from 2.0 to 3.5 % h^-1^ (*P* = 0.01).

**Table 3.** Biometric measurements in pregnant Holstein heifers (n = 11).

| Item | Days of gestation |  |  | P-value |
| --- | --- | --- | --- | --- |
|  | 145 | 200 | 255 |  |
| Body weight, kg | 469.9 ± 20.0<sup>c</sup> | 539.5 ± 20.0<sup>b</sup> | 580.9 ± 20.3<sup>a</sup> | < 0.01 |
| Withers height, m | 1.35 ± 0.01<sup>c</sup> | 1.39 ± 0.01<sup>b</sup> | 1.41 ± 0.01<sup>a</sup> | < 0.01 |
| Croup height, m | 1.41 ± 0.02<sup>c</sup> | 1.44 ± 0.02<sup>b</sup> | 1.47 ± 0.02<sup>a</sup> | < 0.01 |
| Body length, m | 1.21 ± 0.04<sup>b</sup> | 1.48 ± 0.04<sup>a</sup> | 1.49 ± 0.04<sup>a</sup> | < 0.01 |
| Thoracic perimeter, m | 1.89 ± 0.06<sup>b</sup> | 2.03 ± 0.06<sup>a</sup> | 2.10 ± 0.06<sup>a</sup> | < 0.01 |

<sup>a-b</sup>Means within a row with different superscripts differ ( $P < 0.05$ ).

**Table 4.** Digestion kinetics for Holstein heifers in different days of gestation (n = 8).

| Item |  | Days of gestation |  |  | *P*-value |
| --- | --- | --- | --- | --- | --- |
|  |  | 145 | 200 | 255 |  |
| Rates, % h<sup>-1</sup> |  |  |  |  |  |
| Dry matter | *k<sub>i</sub>*<sup>1</sup> | 7.3 ± 0.65 | 8.4 ± 0.65 | 8.3 ± 0.72 | 0.25 |
|  | *k<sub>p</sub>*<sup>2</sup> | 4.5 ± 0.43 | 4.8 ± 0.43 | 4.9 ± 0.47 | 0.57 |
|  | *k<sub>d</sub>*<sup>3</sup> | 2.8 ± 0.36 | 3.6 ± 0.36 | 3.4 ± 0.41 | 0.24 |
| Neutral detergente fiber | *k<sub>i</sub>* | 4.8 ± 0.37 | 4.5 ± 0.37 | 5.0 ± 0.41 | 0.20 |
|  | *k<sub>p</sub>* | 2.2 ± 0.36 | 2.3 ± 0.36 | 1.6 ± 0.42 | 0.22 |
|  | *k<sub>d</sub>* | 2.0 ± 0.26<sup>b</sup> | 2.6 ± 0.26<sup>ab</sup> | 3.5 ± 0.30<sup>a</sup> | 0.01 |
| Rumen pool sizes, kg |  |  |  |  |  |
| Dry matter |  | 4.9 ± 0.28 | 5.0 ± 0.28 | 4.6 ± 0.30 | 0.36 |
| Neutral detergente fiber |  | 2.5 ± 0.16 | 2.2 ± 0.16 | 2.2 ± 0.18 | 0.14 |

<sup>1</sup> $k_i$ = rate of ingestion of feed fractions.
<sup>2</sup> $k_p$ = passage rate of feed fractions.
<sup>3</sup> $k_d$ = digestion rate of feed fractions.

There was no difference (*P* > 0.05) for the ruminal pool of dry matter and neutral detergent fiber among days of gestation, thereby presenting with an average ruminal pool of 3.9 kg for dry matter and 2.3 kg for neutral detergent fiber.

### Ruminal and omasal pH

There was pH interaction (*P* = 0.01) between collection times and days of gestation for ruminal fluid and omasal fluid (*P* < 0.01) (Figs 2 and 3). Values for ruminal pH increased (*P* < 0.01) at 0500 h from 145 to 255 days of gestation, the same pattern was observed at 1700, 2000 and 2300 h (*P* < 0.01) with pH values from 5.5 to 6.11, 5.6 to 5.9 and 5.6 to 6.2, respectively. The omasal pH interaction effect was present at 0500, 0800, 1100 and 1400 h. Values of pH at 0500 increased in 10.8 % from 145 to 200 days of gestation, and the pH values at 0800, 1100 and 1400 h increased 2.6%, 6.1% and 6.0% from 145 to 255 days of gestation, respectively.

**Figure 2.**
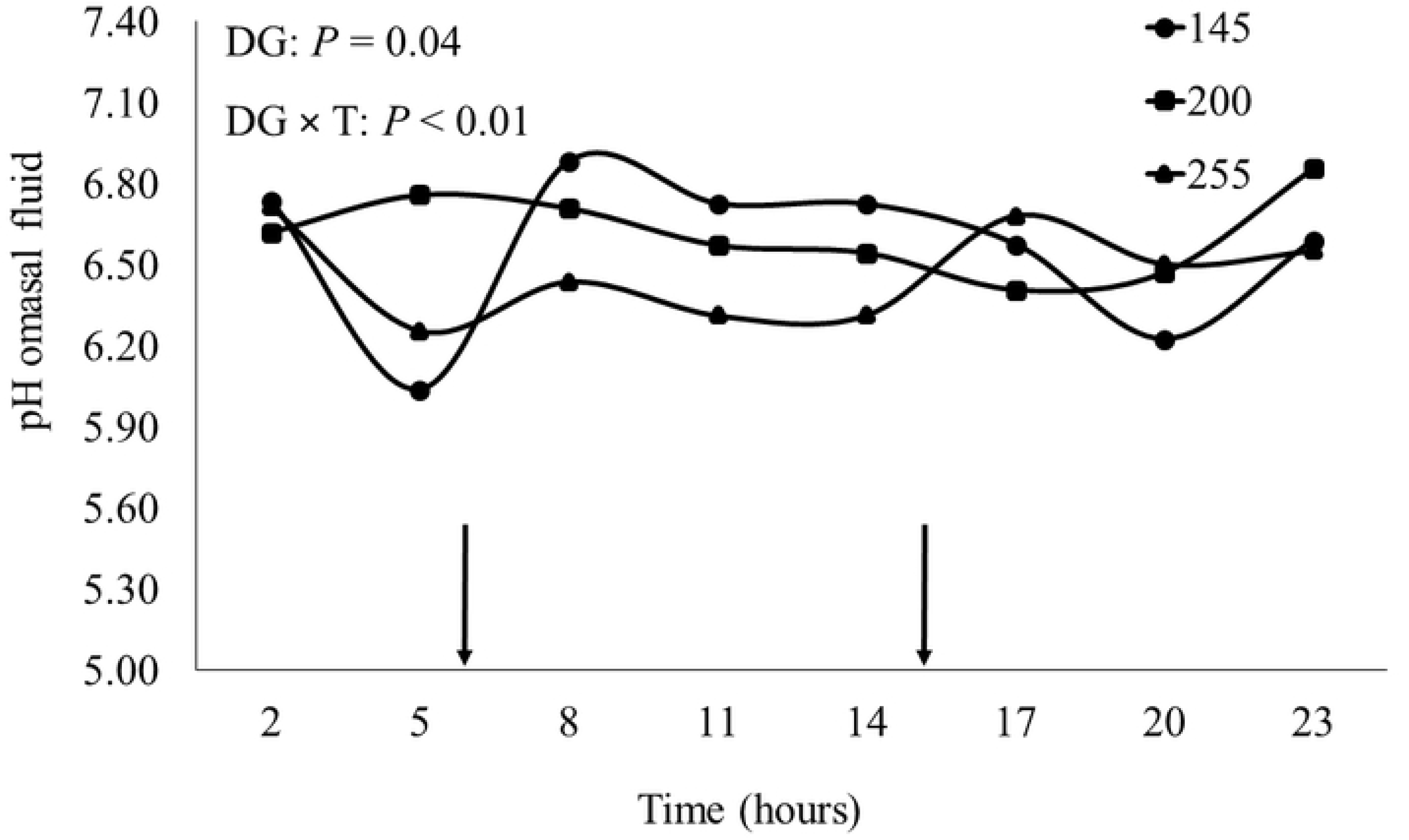
pH of ruminal fluid in different days of gestation of Holstein heifers. Arrows indicate the time of diet delivered at 0700 and 1500 h (number of animals = 8).

**Figure 3.**
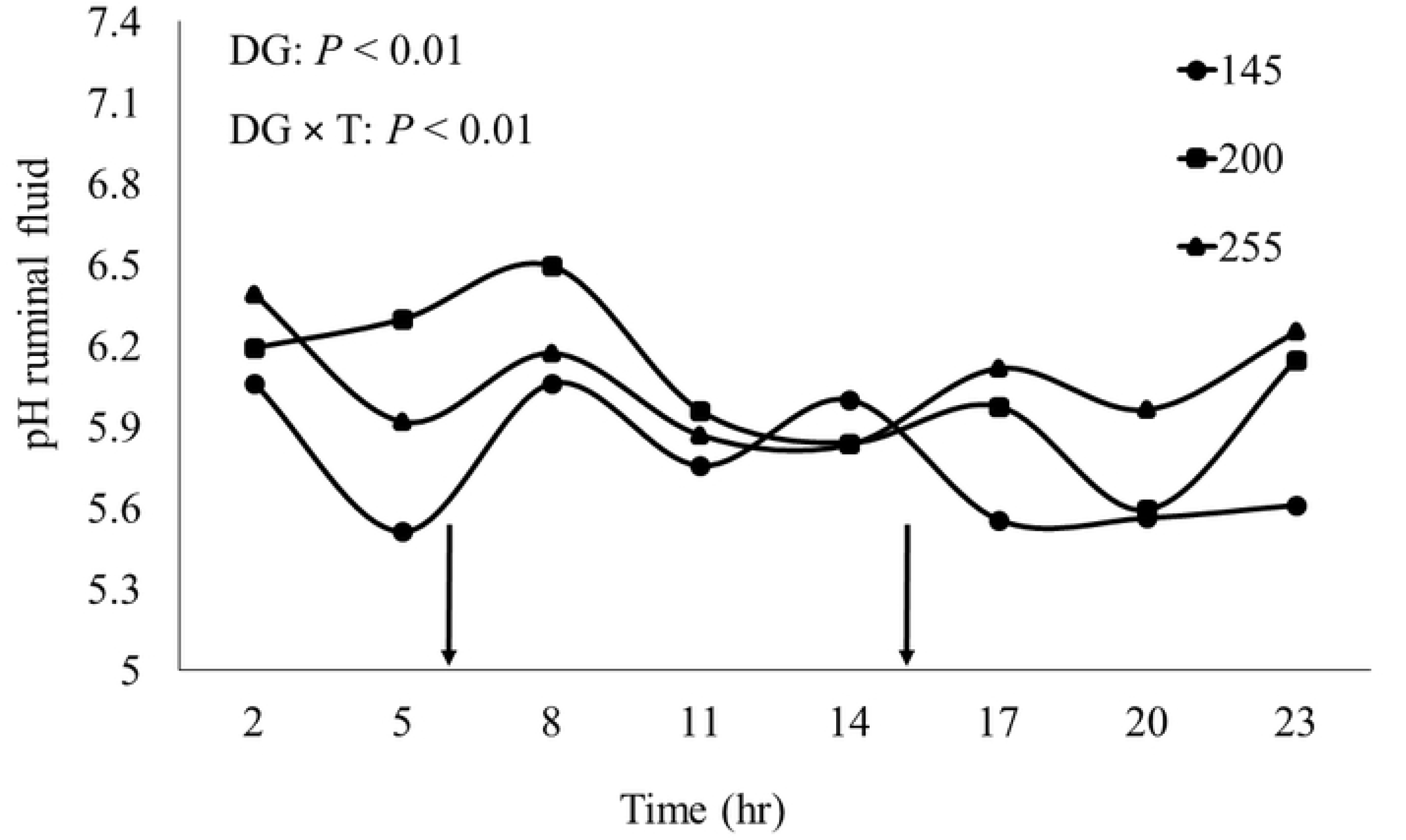
pH of omasal fluid in different days of gestation of Holstein heifers. Arrows indicate the time of diet delivered at 0700 and 1500 h (number of animals = 8).

### Body measurements

The measurements of the teats were not different (*P* > 0.05) among the evaluated periods, however, we observed that the anterior teats were 15.1% larger than the posterior teats (Table 3). The measurements of body weight, withers height, croup height, body length, and thoracic perimeter presented a difference (*P* < 0.01) among periods that were evaluated with an increase of 23.7, 4.4, 4.2, 23.1 and 11.1% respectively, when we compare 145 and 255 days of gestation (Table 3).

### Blood metabolites

Glucose levels decreased (*P* = 0.01) by approximately 6.9% from 145 to 255 days of gestation and the levels of β-hidroxibutirate increased (*P* < 0.01) in the same period by 62.9% (Table 5). The urea and nonesterified fatty acids concentration did not differ (*P* > 0.05) among days of gestation.

**Table 5.** Metabolic parameters (mmol/L) of blood samples for Holstein heifers in different days of gestation (n = 11).

| Item | Days of gestation |  |  | P-value |
| --- | --- | --- | --- | --- |
|  | 145 | 200 | 255 |  |
| Glucose | $4.0 \pm 0.05^a$ | $3.9 \pm 0.05^{ab}$ | $3.8 \pm 0.06^b$ | $< 0.01$ |
| Non-esterified fatty acids | $0.2 \pm 0.04$ | $0.2 \pm 0.04$ | $0.3 \pm 0.05$ | 0.14 |
| Urea | $1.7 \pm 0.12$ | $1.8 \pm 0.12$ | $1.8 \pm 0.14$ | 0.49 |
| β-hydroxybutyrate | $0.3 \pm 0.04^b$ | $0.4 \pm 0.04^a$ | $0.4 \pm 0.04^a$ | $< 0.01$ |

<sup>a-b</sup>Means within a row with different superscripts differ ( $P < 0.05$ ).

## Discussion

There is a lack of information about dry matter intake changes of growing heifers during the last trimester of gestation [9]. To our knowledge this is the first study aiming to evaluate digestive parameters in pregnant Holstein heifers. When comparing dry matter intake (kg/day) at three days of gestation evaluated, we observed an increase of approximately 15.5% between 145 and 200 days of gestation (1.3 kg/day). This might be explained by the fact that heifers are still growing (increasing body measurements) and had an average daily gain around 1.0 kg which is not the case of mature cows, which have a reduction in dry matter intake during pregnancy [12, 24, 25]. Similarly, nutrient intake also showed the same pattern as dry matter in this study.

However, we observed that dry matter intake expressed in % body weight was constant at 145 and 200 days of gestation and had a 16.7% decrease at 255 days of gestation, which corresponds to the 37^th^ week of gestation, approximately. This reduction was also reported by Park [26] (25.6%), and Rotta [12] (22.0%) when evaluating dairy cows. However, the reduction of dry matter intake expresses in % body weight, from dairy cows reported was numerically greater than our study with pregnancy heifers. The reduction of dry matter intake might be influenced by increase or decrease in the concentration of hormones such as estrogen and progesterone, respectively [27], as well as the reduction of rumen physical space at the end of gestation [12], which, according to Forbes [28], is the most limiting factor of consumption. This result implies a greater use of feed by the animal.

According to Rotta [12] and Park [26], with advancing gestation the total-tract dry matter, organic matter, crude protein and neutral detergent fiber digestibility, has shown a reduction, which in part can be explained by the increase of passage rate [29]. Nevertheless, in the present study, we observed that apparent total-tract dry matter, organic matter, crude protein and neutral detergent fiber digestibility increased and no difference was observed for the *k*_*p*_ in the period of gestation evaluated, which explains the non-reduction of total-tract digestibility.

Studies evaluating digestion kinetics during gestation of cows are scare. In our study the rates of gestation, passage and digestion remained constant, with no differences among the days of gestation, excepted for the neutral detergent fiber *k*_*d*_. Comparing multiparous cows [30,31] observed an increase in the rate of passage during the pregnancy, with decrease nutrient digestibility [32]. However, the pattern observed for pregnant dairy cows was not observed in this study. Factors such as physiological, gastrointestinal, and feed changes should be considered to explain response rates in addition to feed characteristics (particle size) [26] and might have influenced the lack of difference along the days of gestation when these variables were assessed.

Further studies should be performed to better elucidate rumen parameters at late gestation in Holstein heifers, so they may be used as the basis for formulating rations to meet the requirements of this category, since the focus in the studies are based on multiparous lactating cows.

Considering the concentration of metabolites in dairy cows, studies using lactating cows have been carried out and limits of several metabolites have been established [33, 34, 35]. However, according to Briscic [36], reference values for heifers at the end of gestation have not been studied. In this study, values for glucose and urea were similar to those found by Briscic [36]: minimum concentration of 2.8 mmol/L and maximum of 4.0 mmol/L for glucose, and for urea the values were between 1.1 mmol/L and 5.8 mmol/L, thus establishing such values as a reference for heifers during late-gestation.

Values of β-hidroxibutirate in the present study reached 0.44 mmol/L, which are 3 times lower than the values utilized as threshold for ketosis [37]. The values found for nonesterified fatty acids reached 0.34 mmol/L. According to Overton [33], levels above 0.5 mmol/L in the prepartum period indicate a greater probability of developing some metabolic disease. Thus, the heifers in the present study with an average daily gain approximately of 1.0 kg are unlikely to present metabolic problems due to the low levels of β-hidroxibutirate and nonesterified fatty acids.

## Conclusions

Holstein heifers during the last trimester of pregnancy demonstrate an increase in the intake and apparent total-tract digestibility of dry matter, organic matter, and crude protein from 200 days of gestation. Also, there was a reduction of dry matter intake expressed as a percentage of body weight at the end of gestation by 16.7%, reaching 1.49% body weight at 250 days of gestation. The digestion kinetics remained constant. Studies evaluating such factors are scarce in the literature, so further research is still needed to consolidate such results.

## Acknowledgments

The authors thank the Instituto Nacional de Ciências e Tecnologia de Ciência Animal (INCT-CA), Fundação de Amparo à Pesquisa do Estado de Minas Gerais (FAPEMIG), Coordenação de Aperfeiçoamento de Pessoal de Nível Superior (CAPES) and Conselho Nacional de Desenvolvimento Científico e Tecnológico (CNPq, Brazil) for financial support.

